# Combining multi-omics and drug perturbation profiles to identify novel treatments that improve disease phenotypes in spinal muscular atrophy

**DOI:** 10.1101/2019.12.17.879353

**Authors:** Katharina E. Meijboom, Viola Volpato, Jimena Monzón-Sandoval, Joseph M. Hoolachan, Suzan M. Hammond, Frank Abendroth, Olivier Gerrit de Jong, Gareth Hazell, Nina Ahlskog, Matthew J.A. Wood, Caleb Webber, Melissa Bowerman

## Abstract

Spinal muscular atrophy (SMA) is a neuromuscular disorder caused by loss of survival motor neuron (SMN) protein. While SMN restoration therapies are beneficial, they are not a cure. We aimed to identify novel treatments to alleviate muscle pathology combining transcriptomics, proteomics and perturbational datasets. This revealed potential drug candidates for repurposing in SMA. One of the lead candidates, harmine, was further investigated in cell and animal models, improving multiple disease phenotypes, including SMN expression and lifespan. Our work highlights the potential of multiple, parallel data driven approaches for development of novel treatments for use in combination with SMN restoration therapies.

## INTRODUCTION

Spinal muscular atrophy (SMA) is an autosomal recessive neuromuscular disorder ^1^ and the leading genetic cause of infant mortality ^2^. The major pathological components of the disease are the selective loss of spinal cord alpha motor neurons, progressive muscle denervation ^3^ and skeletal muscle atrophy ^4^. SMA is caused by mutations in the *survival motor neuron 1* (*SMN1*) gene ^5^. SMN protein is ubiquitously expressed and complete loss is lethal ^6^. However, humans have a near-identical centromeric copy of the *SMN1* gene, termed *SMN2*, in which a single nucleotide change (C to T) in exon 7 ^7^ results in the exclusion of exon 7 from ∼90% of the mature transcript ^8^. The resulting protein is unstable and gets rapidly degraded ^9^. Patients can have a varying number of *SMN2* copies, which correlates with disease severity as each *SMN2* copy retains the ability to produce ∼10% of functional full-length (FL) protein ^10, 11^.

The first SMN restoration treatments, Spinraza™ and Zolgensma™, have recently been approved by the Food and Drugs Administration (FDA) and the European Medicine Agency (EMA). Spinraza™ ^12^ is an antisense oligonucleotide (ASO) that promotes *SMN2* exon 7 inclusion ^13^ and is administered by lumbar puncture while Zolgensma™ delivers *SMN1* cDNA via an adeno-associated virus 9 ^14^ and is administered intravenously. Additional systemically delivered and SMN targeting small molecules are currently being explored in clinical trials such as risdiplam ^15, 16^. While these treatments have changed the SMA therapeutic landscape, they unfortunately fall short of representing a cure ^17–20^. There is therefore a present need for SMN-independent therapies that could be used in combination with SMN restoration treatments to provide a longer-lasting and more effective therapeutic management of SMA pathology in patients ^17–19^.

Skeletal muscle pathology is a clear contributor to SMA disease manifestation and progression and improving muscle health could have significant benefits for patients ^21^. Here, we used an in-depth, parallel approach combining proteomics, transcriptomics and the drug pertubational dataset Connectivity Map (CMap) ^22, 23^ to identify differentially expressed (DE) genes and proteins in skeletal muscle of the severe Taiwanese *Smn^-/-^;SMN2* SMA mice ^24^ that could potentially be restored by known and available pharmacological compounds. This strategy uncovered several potential therapeutic candidates, including harmine, which was further evaluated in cell and animal models, showing an ability to restore molecular networks and improve several disease phenotypes, including SMN expression and lifespan. Our study highlights the tremendous potential of intersecting disease multi-omics with drug perturbational responses to identify therapeutic compounds capable of modulating dysfunctional cellular networks to ameliorate SMA phenotypes.

## RESULTS

### Early restoration of Smn in SMA mice restores muscle protein and transcript expression

We first set out to determine the effect of early SMN restoration on the proteomic and transcriptomic profiles of SMA skeletal muscle, with the intent to design therapeutic strategies against the genes and proteins that remain unchanged. To do so, the severe Taiwanese *Smn^-/-^;SMN2* SMA mouse model ^24^ received a facial intravenous (IV) injection at post-natal day (P) 0 and P2 of the previously described Pip6a-PMO or Pip6a-scrambled pharmacological compounds (10 μg/g) ^25, 26^. Pip6a is a cell-penetrating peptide (CPP) either conjugated to an *SMN2* exon 7 inclusion-promoting ASO (PMO) or a scrambled ASO ^25, 26^. We harvested the *tibialis anterior* (TA) from P2 (pre-symptomatic) untreated *Smn^-/-^;SMN2* and wild type (WT) mice, P7 (symptomatic) untreated *Smn^-/-^;SMN2* and WT mice and P7 Pip6a-scrambled- and Pip6a-PMO-treated *Smn^-/-^;SMN2* mice. TAs were then cut in two, whereby one half was used for transcriptomics and the other for proteomics. qPCR analysis of the ratio of FL *SMN2* over total *SMN2* confirms a significant increase in FL SMN2 expression in P7 Pip6a-PMO-treated *Smn^-/-^;SMN2* mice compared to age-matched untreated and Pip6a-scrambled-treated *Smn^-/-^;SMN2* mice (Fig. 1a).

**Figure 1.**
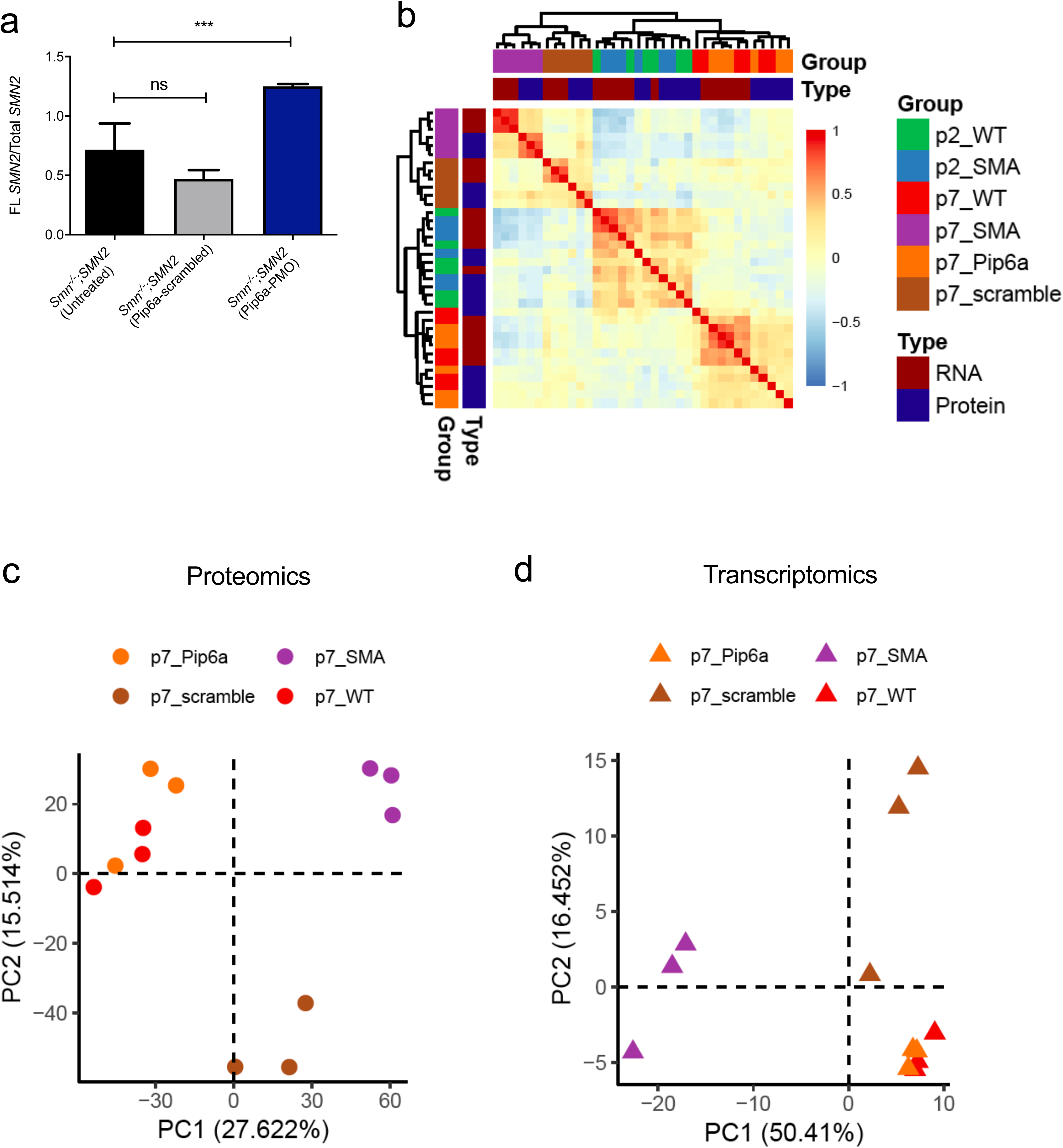
Restoration of protein and transcript expression in skeletal muscle of SMA mice following early SMN restoration treatment. *Smn^-/-^;SMN2* mice received a facial intravenous injection at postnatal day (P) 0 and P2 of Pip6a-scrambled or Pip6a-PMO (10 μg/g). The *tibialis anterior* was harvested from P2 untreated *Smn^-/-^;SMN2* and WT mice, P7 untreated, Pip6a-scrambled-treated and Pip6a-PMO-treated *Smn^-/-^;SMN2* mice and P7 untreated WT mice. **a.** Comparison of the ratio of full length (FL) *SMN2* over total *SMN2* quantified by qPCR between P7 untreated Pip6a-scrambled- and Pip6a-PMO-treated *Smn^-/-^;SMN2* mice. Data are mean ± s.d., n = 4 animals per experimental group, one-way ANOVA, ns = not significant, \*\*\**p*<0.001. **b.** Heatmap shows the similarity between transcriptomic and proteomic expression profiles measured by the Pearson correlation between each pair of samples (after the removal of the first principal component). **c.** First two principal components constructed from proteomic profiles mice discriminate Smn-/-;SMN2 mice from WT at P7. Notably, only mice treated with Pip6a-PMO cluster along with WT mice. **d**. First two principal components based on transcriptional profiles of P7 mice show similar clustering of Pip6a-PMO cluster and WT mice.

Despite differences between transcriptomic and proteomic methodologies highlighted by hierarchical clustering and combined Principal Component Analysis (PCA) (Supplementary Fig. 1), we were able to find clear separation of experimental groups and agreement between transcriptomic and proteomic profiles once the variance attributed to the differences in methodologies was removed (Fig. 1b). At P7, we observed clear separation of *Smn^-/-^;SMN2^-^* and WT samples, where only P7 Pip6a-PMO treated *Smn^-/-^; SMN2^-^* mice clustered with WT. P2 *Smn^-/-^;SMN2* and WT samples cluster together suggesting that overt disease cannot be detected in omics readouts at this early stage (Fig. 1b, Supplementary Fig. 2). In the PCA of P7 samples only (Fig. 1c for proteomics and Fig. 1d for transcriptomics), we noted clustering of P7 Pip6a-PMO-treated *Smn^-/-^;SMN2* mice with untreated P7 WT animals, implying full restoration to normal phenotypes. Surprisingly, we also detected segregation of Pip6a-scrambled-treated samples at both transcriptomics and proteomics levels, revealing that presence of the CPP itself impacts transcription and translation (Fig.1 c,d). Importantly, both the combined and separate analysis of transcriptomic and proteomic data allowed us to identify a robust SMA disease signature in muscle and a Pip6a-PMO treatment efficacy signature. Indeed, identification of differentially expressed genes and proteins reveals that early induction of FL *SMN* expression by Pip6a-PMO normalizes the expression of all transcripts and all but 11 proteins in the TA of *Smn^-/-^;SMN2* mice (Table 1, Supplementary Table 1). Of note, one of the proteins that remained significantly downregulated is Smn itself (Supplementary Table 1).

**Table 1.** Number of differentially expressed (DE) transcripts and proteins between experimental groups.

Our in-depth molecular profiling thus demonstrates for the first time that increasing FL *SMN2* in neonatal SMA mice quasi-completely normalizes muscle transcripts and proteins, highlighting at the molecular level the potential treatment benefits arising from early intervention.

### CMap perturbational profiles identify potential novel non-SMN treatments

We used the transcriptomic and proteomic profiles of the *Smn^-/-^;SMN2* mice treated with Pip6a-PMO to find drugs that induced similar transcriptional patterns using the Connectivity Map (CMap) resource ^27, 28^. For this, we obtained a cleaned and reversed disease signature for both transcriptomics and proteomics data by excluding the genes and proteins restored by Pip6a-scrambled (Pip6a-scrambled-treated *Smn^-/-^;SMN2 vs* untreated WT) from the overlap between disease (untreated *Smn^-/-^;SMN2* vs untreated WT) and Pip6a-PMMO (Pip6a-PMO treated *Smn^-/-^;SMN2 vs* untreated *Smn^-/-^;SMN2*) (Fig. 2a). Although these cleaned sets of transcripts and proteins did not show high overlap between omics data types (Fig. 2b), we found similarity at the level of enriched pathways (Fig. 2c). Individual pathway analysis for the disease and the Pip6a-PMO treatment are compiled in Supplementary file 1. The top 10 pharmacological compounds from CMap that showed a reversed pattern of expression for the disease signature, and similar expression patterns to those observed with Pip6a-PMO treatment are listed in Table 2. Importantly, a subset of these drugs, namely salbutamol^29^ and alsterpaullone^30^, have already been considered for SMA treatment, highlighting the capability of this analytic approach to identify relevant therapeutic options for SMA.

**Figure 2.**
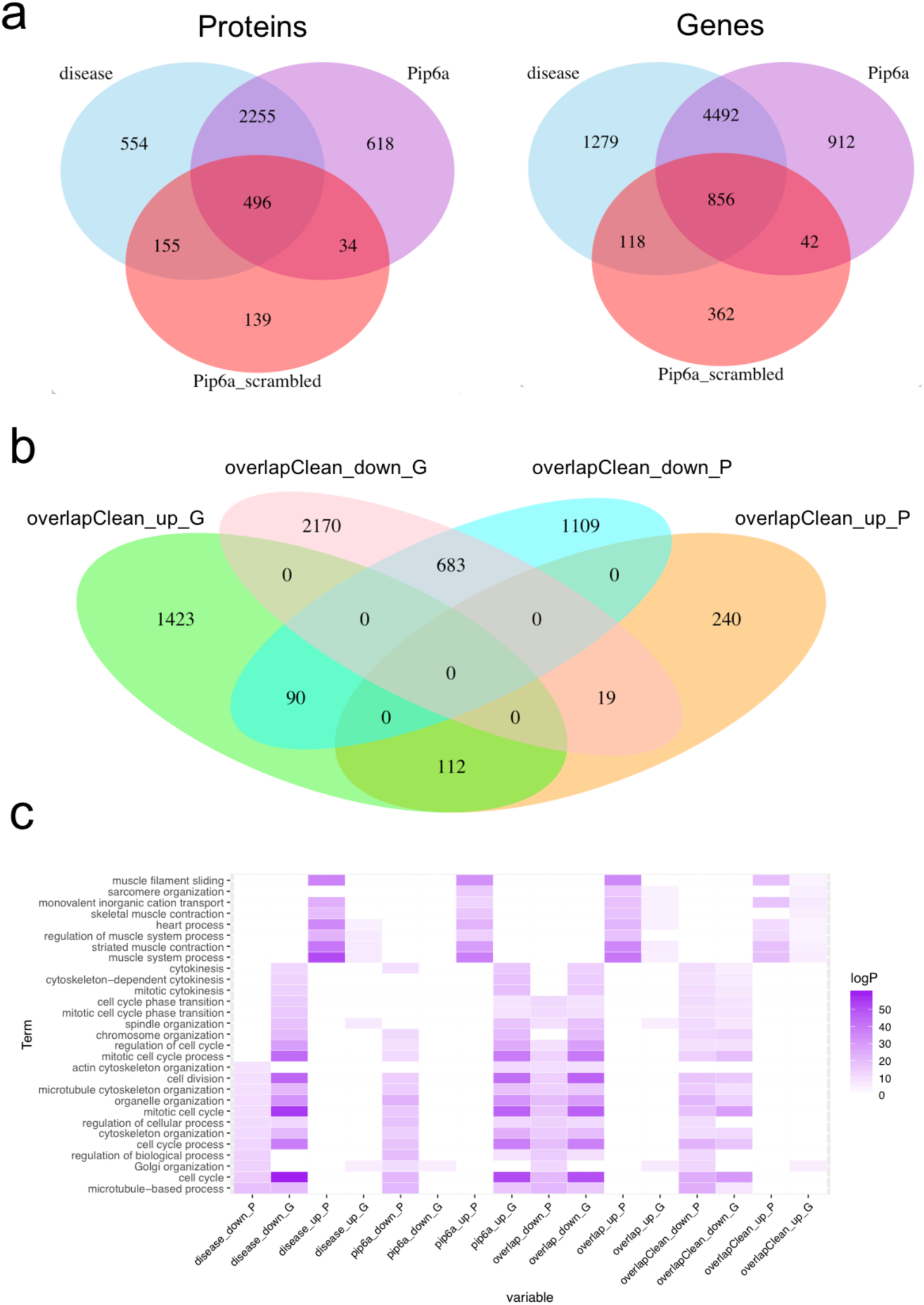
Identification of disease signal reversed by treatment with Pip6a at both proteomics and transcriptomics level. **a**. We retained the proteins (left) and genes (right) that were differentially expressed between untreated *Smn^-/-^;SMN2* and WT (disease), reversed by treatment with Pip6a and not differentially expressed between scrambled-Pip6a treated *Smn^-/-^;SMN2* and WT. We called these as cleaned signatures. **b**. Overlap of the cleaned signatures between proteins and genes. Although the overlap was not high, we detected higher overlaps for genes and proteins with the same directionality (both up or both down). **c**. Enriched GO Biological Processes terms that showed similarity across comparisons. The greatest similarity between genes and proteins was detected for the cleaned signatures.

**Table 2.** Top 10 top pharmacological compounds identified by CMap analysis based on three expression signatures for both the transcriptomic data and proteomic data.

To further validate our approach, we chose to evaluate the potential of harmine (chemically akin to harmol), a drug identified by its CMap profile but not previously evaluated for SMA, which was present in several proteomic and transcriptomic signatures (Table 2). Harmine is an alkaloid isolated from the seeds of Peganum harmala, traditionally used for ritual and medicinal preparations ^31, 32^. Harmine has also demonstrated therapeutic benefits ^33^ in animal models of the motor neuron disease amyotrophic lateral sclerosis (ALS) ^34^ and the muscle disorder myotonic dystrophy type 1 (DM1) ^35^.

We firstly validated the genes and proteins predicted to be dysregulated by the transcriptomics and proteomics data and to be normalized by harmine through the CMap analysis. We indeed confirm by qPCR analysis that the genes *Snrnp27*, *Gls*, *Aspm* and *Mcm2* are significantly downregulated while *Clpx*, *Ppm1b, Tob2* and *Cdkn1a* are significantly upregulated in the TA of P7 *Smn^-/-^;SMN2* mice compared to WT animals (Fig. 3a). We then evaluated the ability of harmine to impact the expression of these genes by treating C2C12 myoblasts, NSC-34 neuronal-like cells, SMA patient fibroblasts and control fibroblasts with 25, 35 and 50 µM of the drug for 48 hours. We find that harmine demonstrates its predicted activity in a cell- and dose-dependent manner (Fig. 3b). Of note, harmine also displayed inhibitory effects on proliferation and viability at the higher doses in C2C12s and NSC-34s (Supplementary Fig. 3). Finally, we investigated the influence of harmine on SMN expression and observe a significant increased *Smn* expression in C2C12s and NSC34s at several doses (Fig. 3c). Interestingly, we also find a significant upregulation of FL *SMN2* in SMA patient fibroblasts without any changes in total *SMN2* (Fig. 3d).

**Figure. 3.**
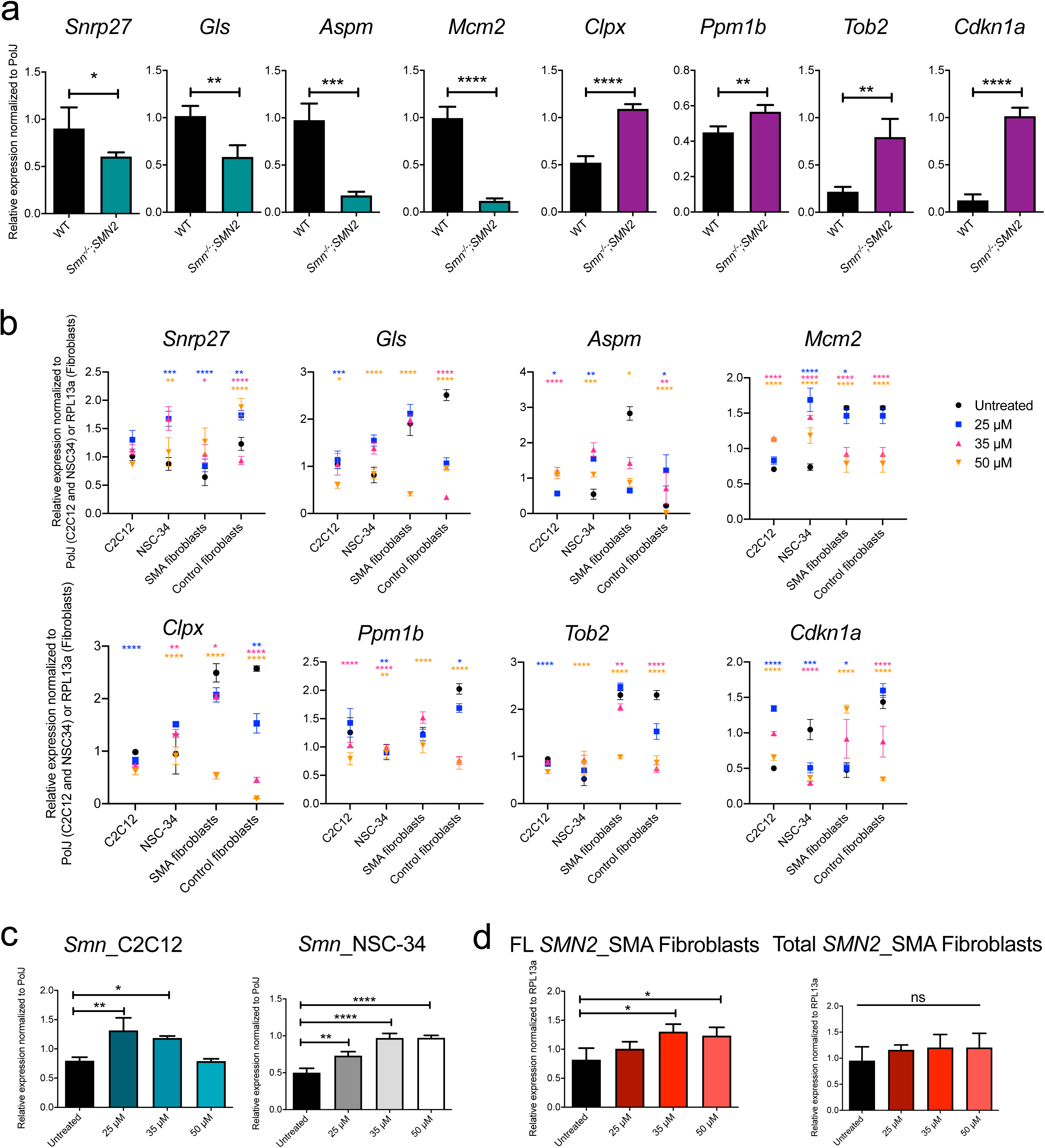
Harmine, as predicted by CMap analyses, is able to reverse the expression of genes differentially expressed in SMA muscle in several cellular models. **a.** qPCR analysis of genes predicted to be significantly downregulated (*Snrnp27*, *Gl*s, *Aspm* and *Mcm2*) and upregulated (*Clpx*, *Ppm1b*, *Tob2* and *Cdkn1a*) in the TA of untreated P7 SMA *Smn^-/-^;SMN2* and WT mice. Data are mean ± s.d., n = 4 animals per experimental group, *t* test, *p* = 0.041 (*Snrnp27*), *p* = 0.0019 (*Gl*s), *p* = 0.0001 (*Aspm*), *p*<0.0001 (*Mcm2)*, *p*<0.0001 (*Clpx*), *p* = 0.0076 (*Ppm1b*), *p* = 0.0012 (*Tob2*), *p*<0.0001 (*Cdkn1a*). **b.** C2C12s, NSC34s, SMA fibroblasts and control fibroblasts were treated with 25, 35 of 50 µM of harmine for 48 hours. Expression of *Snrnp27*, *Gl*s, *Aspm*, *Mcm2*, *Clpx*, *Ppm1b*, *Tob2* and *Cdkn1a* was assessed by qPCR and compared to untreated cells. Data are mean ± s.d., n = 3 independent wells, two-way ANOVA, \**p*<0.05, \*\**p*<0.01, \*\*\**p*<0.001, \*\*\*\**p*<0.0001. **c.** C2C12s and NSC34s were treated with 25, 35 of 50 µM of harmine for 48 hours. Expression of *Smn* was assessed by qPCR and compared to untreated cells. Data are mean ± s.d., n = 3 independent wells, one-way ANOVA, \**p*<0.05, \*\**p*<0.01, \*\*\*\**p*<0.0001. **d.** SMA fibroblasts were treated with 25, 35 of 50 µM of harmine for 48 hours. Expression of FL *SMN2* and total *SMN2* was assessed by qPCR and compared to untreated cells. Data are mean ± s.d., n = 3 independent wells, one-way ANOVA, ns = not significant, \**p*<0.05.

Thus, our strategy of combining transcriptomics, proteomics and drug perturbational datasets has resulted in the generation of a list of several drugs with the potential to restore muscle health in SMA. Importantly, selecting harmine for additional proof-of-concept investigations, highlights the strength of this approach.

### Administration of harmine to SMA mice ameliorates disease phenotypes

To further evaluate the therapeutic effects of harmine *in vivo*, we administered it daily to *Smn^-/-^;SMN2* mice and *Smn^+/-^;SMN2* control littermates by gavage (10 mg/kg diluted in 0.9% saline) starting at P0. We first evaluated the effects of harmine on the expression of *Snrnp27*, *Gls*, *Aspm*, *Mcm2*, *Clpx*, *Ppm1b, Tob2* and *Cdkn1a* in muscle (triceps) of P7 untreated and harmine-treated *Smn^-/-^;SMN2* and *Smn^+/-^;SMN2* mice (Fig. 4a). *In vivo*, harmine only impacted the expressions of *Snrnp27* and *Tob2* in *Smn^-/-^;SMN2* mice, towards normalized levels (Fig. 4a). Similar to that observed *in vitro*, harmine administration significantly increased FL *SMN2* expression in *Smn^-/-^;SMN2* mice but not total *SMN2* (Fig. 4b). Total SMN protein levels were also not affected by harmine (Fig. 4c).

**Figure 4.**
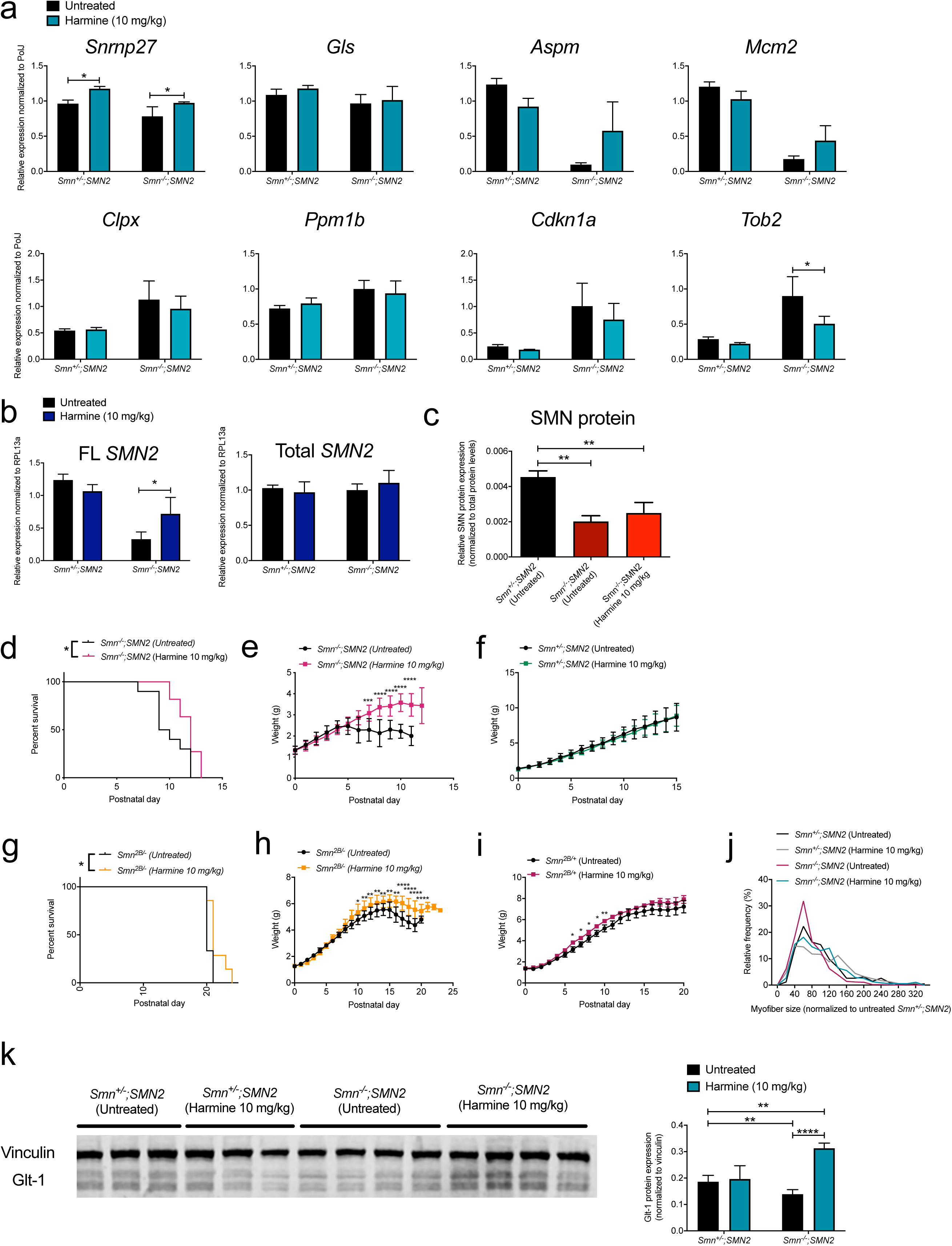
Administration of harmine to SMA mice ameliorates several disease phenotypes. All treated animals received a daily dose of harmine (10 mg/kg, diluted in 0.9% saline) by gavage starting at postnatal day (P) 0. **a.** qPCR analysis of *Snrnp27*, *Gl*s, *Aspm*, *Mcm2*, *Clpx*, *Ppm1b*, *Tob2* and *Cdkn1a* in triceps of P7 untreated and harmine-treated *Smn^-/-^;SMN2* mice and *Smn^+/-^;SMN2* control littermates. Data are mean ± s.d., n = 4 animals per experimental group except for harmine-treated *Smn^+/-^;SMN2* where n = 3, two-way ANOVA, \**p*<0.05. **b.** qPCR analysis of FL *SMN2* and total *SMN2* in triceps of P7 untreated and harmine-treated *Smn^-/-^;SMN2* mice and *Smn^+/-^;SMN2* control littermates. Data are mean ± s.d., n = 4 animals per experimental group except for harmine-treated *Smn^+/-^;SMN2* where n = 3, two-way ANOVA, \**p*<0.05. **c.** Western blot analysis of SMN protein in triceps of P7 untreated *Smn^+/-^;SMN2* control littermates and P7 harmine-treated and untreated *Smn^-/-^;SMN2* mice and. Data are mean ± s.d., n = 3 animals per experimental group, two-way ANOVA, \*\**p*<0.01. **d.** Survival curves of untreated and harmine-treated *Smn^-/-^;SMN2* mice. Data are Kaplan Meier survival curve, n = 10 for untreated *Smn^-/-^;SMN2* mice, n = 11 for harmine-treated *Smn^-/-^;SMN2* mice, Log-rank (Mantel-Cox) test, \**p* = 0.0211. **e.** Daily weights of untreated and harmine-treated *Smn^-/-^;SMN2* mice. Data are mean ± s.d., n = 10 for untreated *Smn^-/-^;SMN2* mice, n = 11 for harmine-treated *Smn^-/-^;SMN2* mice, two-way ANOVA, \*\*\**p*<0.001, \*\*\*\**p*<0.0001. **f.** Daily weights of untreated and harmine-treated *Smn^+/-^;SMN2* mice. Data are mean ± s.d., n = 13 for untreated *Smn^+/-^;SMN2* mice, n = 15 for harmine-treated *Smn^+/-^;SMN2* mice, two-way ANOVA. **g.** Survival curves of untreated and harmine-treated *Smn^2B/−^* mice. Data are Kaplan Meier survival curve, n = 9 for untreated *Smn^2B/−^* mice, n = 7 for harmine-treated *Smn^2B/−^* mice, Log-rank (Mantel-Cox) test, \**p* = 0.0221. **h.** Daily weights of untreated and harmine-treated *Smn^2B/−^* mice. Data are mean ± s.d., n = 9 for untreated *Smn^2B/−^* mice, n = 7 for harmine-treated *Smn^2B/−^* mice, two-way ANOVA, \**p*<0.05, \*\**p*<0.01, \*\*\*\**p*<0.0001. **i.** Daily weights of untreated and harmine-treated *Smn^2B/+^* mice. Data are mean ± s.d., n = 13 for untreated *Smn^2B/+^* mice, n = 8 for harmine-treated *Smn^2B/−^* mice, two-way ANOVA, \**p*<0.05, \*\**p*<0.01. **j.** Relative frequency of myofiber sizes in P7 untreated and harmine-treated *Smn^-/-^;SMN2* and *Smn^+/-^;SMN2* mice. Data are percentages, n = 3 animals per experimental group and >400 myofibers per experimental group. **k.** Representative western blots and quantification of Glt-1/vinculin expression in the spinal cord of P7 untreated and harmine-treated *Smn^-/-^;SMN2* and *Smn^+/-^;SMN2* mice. Data are mean ± s.d., n = 3 for untreated and harmine-treated *Smn^+/-^; SMN2* mice, n = 4 for untreated and harmine-treated *Smn^-/-^;SMN2* mice, two-way ANOVA, \*\**p*<0.01, \*\*\**p*<0.001.

We next assessed the effect of harmine upon disease progression and find a significant increase in survival of harmine-treated *Smn^-/-^;SMN2* mice compared to untreated *Smn^-/-^; SMN2* animals (Fig. 4d). Harmine administration also improved weights of treated *Smn^-/-^;SMN2* mice compared to untreated *Smn^-/-^;SMN2* animals (Fig. 4e). Harmine did not impact the weights of *Smn^+/-^;SMN2* control littermates (Fig. 4f). An intermediate SMA mouse model, termed *Smn^2B/-^* ^36^, was also treated with harmine. Harmine administration to *Smn^2B/−^* mice similarly resulted in a significant increase in survival compared to untreated *Smn^2B/−^* animals (Fig. 4g), albeit to a lesser extent, most likely due to the fact that the treated animals developed tremors and needed to be culled. Tremors have indeed been reported in animal studies of long-term harmine administration ^37–39^. Nevertheless, harmine significantly increased the weights of treated *Smn^2B/−^* mice compared to untreated *Smn^2B/−^* animals (Fig. 4h). Interestingly, harmine also had a small but significant impact on the weights of treated *Smn^2B/+^* control littermates compared to untreated *Smn^2B/+^* animals (Fig. 4i).

Given that harmine was chosen to target molecular effectors in muscle, we measured the myofiber size in the TAs from P7 untreated and harmine-treated *Smn^-/-^;SMN2* and *Smn^+/-^;SMN2* mice. We observe an increased proportion of larger myofibers in harmine-treated *Smn^-/-^;SMN2* mice compared to untreated *Smn^-/-^;SMN2* animals (Fig. 4j).

Finally, harmine has also been been reported to increase the expression of the neuroprotective glutamate transporter 1 (GLT-1) ^40, 41^ and thus, we assessed GLT-1 protein levels in P7 spinal cords from untreated and harmine-treated *Smn^-/-^;SMN2* and *Smn^+/-^;SMN2* mice. We find that GLT-1 levels are significantly lower in untreated *Smn^-/-^; SMN2* mice compared to untreated *Smn^+/-^;SMN2* animals and that harmine administration significantly increases GLT-1 expression in treated *Smn^-/-^;SMN2* mice (Fig. 4k).

We thus demonstrate that treating SMA mice with harmine significantly improves multiple molecular and pathological phenotypes in both skeletal muscle and the spinal cord.

### Harmine administration restores gene expression in muscle of SMA mice

To systematically explore the effects of harmine in SMA muscle, we performed RNA-sequencing (RNA-Seq) on TAs from P7 untreated and harmine-treated *Smn^-/-^;SMN2* and WT mice. A total of 15,523 protein coding genes were identified across all samples. We find that harmine significantly reduces the number of differentially expressed genes in *Smn^-/-^;SMN2* when compared to WT animals (Fig. 5a). Interestingly, harmine treatment in WT animals influences the expression of significantly fewer genes than in *Smn^-/-^;SMN2* mice (Fig. 5b). Finally GO analysis with the number of fully (1038) and partially restored (574) genes identifies several pathways that are positively impacted by harmine in SMA muscle (Fig. 5c), many of which have been previously implicated in SMA pathology such as glucose metabolism ^42^.

**Figure 5.**
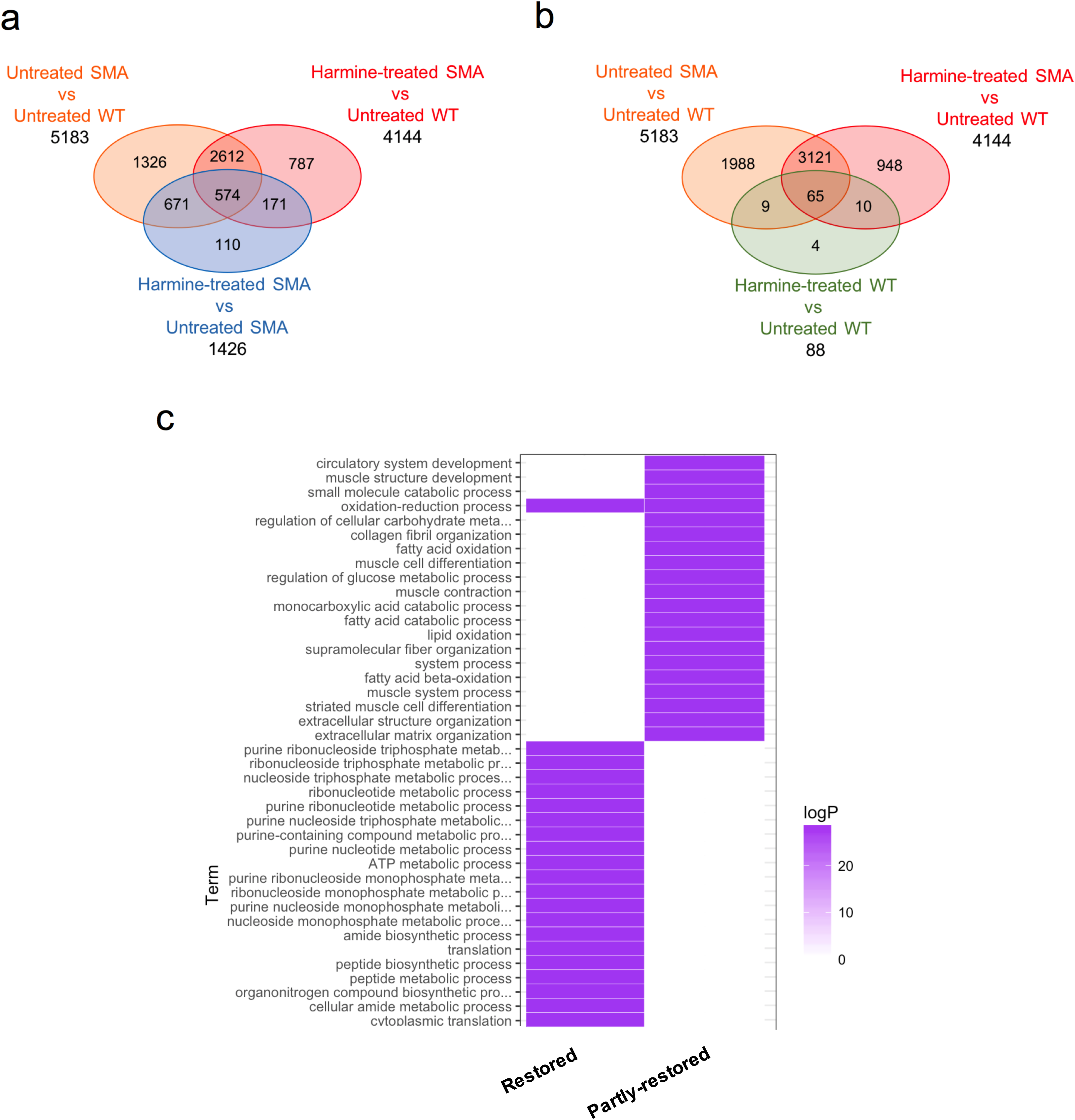
RNA sequencing and pathway analysis reveals full rescue of 20% of dysregulated genes in SMA muscle following harmine administration. All treated animals received a daily dose of harmine (10 mg/kg, diluted in 0.9% saline) by gavage starting at postnatal day (P) 0. TAs were harvested at P7 from untreated and harmine-treated *Smn^-/-^;SMN2* mice and WT animals and processed for RNA sequencing. **a.** Venn diagram representation of the differentially expressed genes based on the negative binomial distribution (DESeq2) in untreated *Smn^-/-^;SMN2* mice vs WT mice, harmine-treated *Smn^-/-^;SMN2* mice vs WT mice and untreated *Smn^-/-^;SMN2* mice vs harmine-treated *Smn^-/-^;SMN2* mice. **b.** Venn diagram representation of the differentially expressed genes based on the negative binomial distribution (DESeq2) in untreated *Smn^-/-^;SMN2* mice vs WT mice, harmine-treated *Smn^-/-^;SMN2* mice vs WT mice and untreated WT mice vs harmine-treated WT mice. **c.** Gene Ontology (GO)/molecular function for fully and partly restored genes in muscle of harmine-treated *Smn^-/-^;SMN2* mice.

Our RNA-Seq analysis therefore supports our earlier prediction that harmine could reverse some of the molecular pathologies in SMA muscle.

### Harmine restores multiple, but not all, molecular networks disturbed in muscle of Smn^-/-^;SMN2 mice

To assess the restorative effects of harmine, we built a gene functional network from the top 500 differentially expressed genes using functional relationships defined by a phenotypic linkage network that links genes together that are likely to influence similar phenotypes ^43^. Louvain clustering of this network identified six modules of interconnected genes disturbed in *Smn^-/-^;SMN2* mice muscle, of which four (M1, M2, M4 and M5) out of six were partially restored by harmine treatment (Fig. 6a). Enrichment analysis in mouse phenotypes highlighted several pathways known to be involved in the pathology such as lipid and glucose metabolism, muscle fiber morphology and contraction (Fig. 6b) providing a molecular explanation for the observed phenotypes in harmine-treated SMA mice and a similarity to the pathways associated with Pip6a-PMO treatment (Fig. 2c). Through Ingenuity Pathway Analysis (IPA), we identified upstream regulators of the restored gene modules (Fig. 6c). The expression of a subset of these upstream regulators was evaluated by qPCR in muscle (triceps) of untreated and harmine-treated *Smn^-/-^;SMN2* and *Smn^+/-^;SMN2* healthy littermates, based on their relevance to muscle health and SMA pathology. We find that in control *Smn^+/-^;SMN2* control animals, harmine significantly impacted the expression of the *dual specificity tyrosine-phosphorylation-regulated kinase 1A* (*Dyrk1a*) and myogenic differentiation 1 (*Myod1*) genes (Fig. 7a). In *Smn^-/-^;SMN2* mice however, harmine only affected the expression of *Myod1* and in the opposite direction (Fig. 7b). To determine if these effects were muscle-specific, we evaluated the expression of *Dyrk1a* and *Myod1* in differentiated WT and siRNA *Smn*-depleted C2C12 cells that were either untreated or exposed to 25 or 50 μM of harmine for 48 hours. Interestingly, we find that harmine significantly reduces *Dyrk1a* and *Myod1* expression in differentiated WT C2C12s without any effect in *Smn*-depleted cells (Fig. 7c,d), revealing differential *in vivo* and *in vitro* effects of harmine. It is therefore evident that additional mechanistic investigations are required to understand the specific and direct benefits of harmine in SMA muscle.

**Figure 6.**
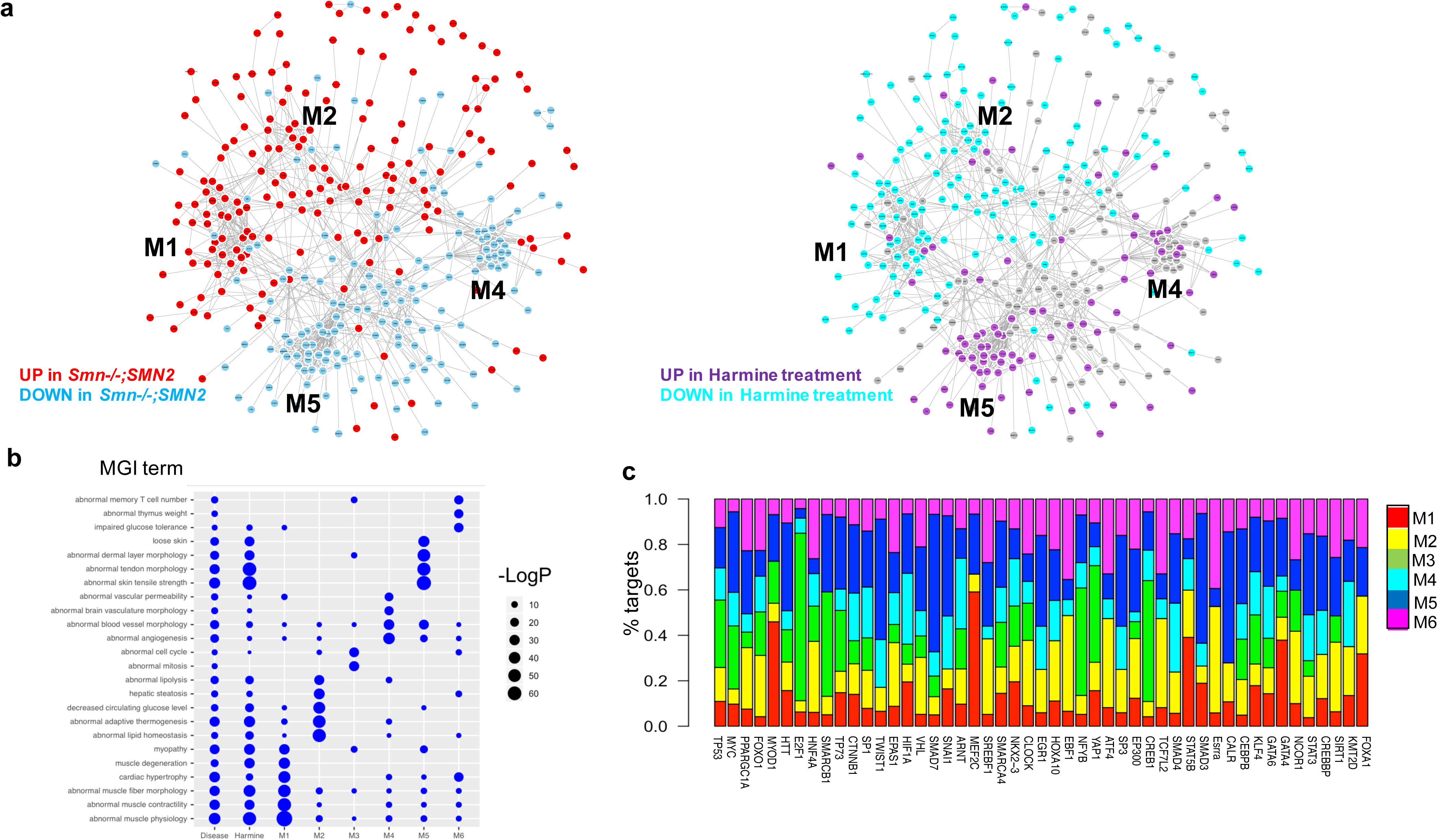
Identification of molecular effectors involved in harmine activity in SMA muscle. **a**. Gene functional network built on the top 500 differentially expressed genes, colored by WT vs SMA (left) and by SMA vs harmine-treated (right). **b**. Top MGI enriched phenotypes for the six identified modules in the network in a. **c**. Proportions of target genes within each of the six modules that are predicted to be regulated by the identified upstream regulators by IPA.

**Figure 7.**
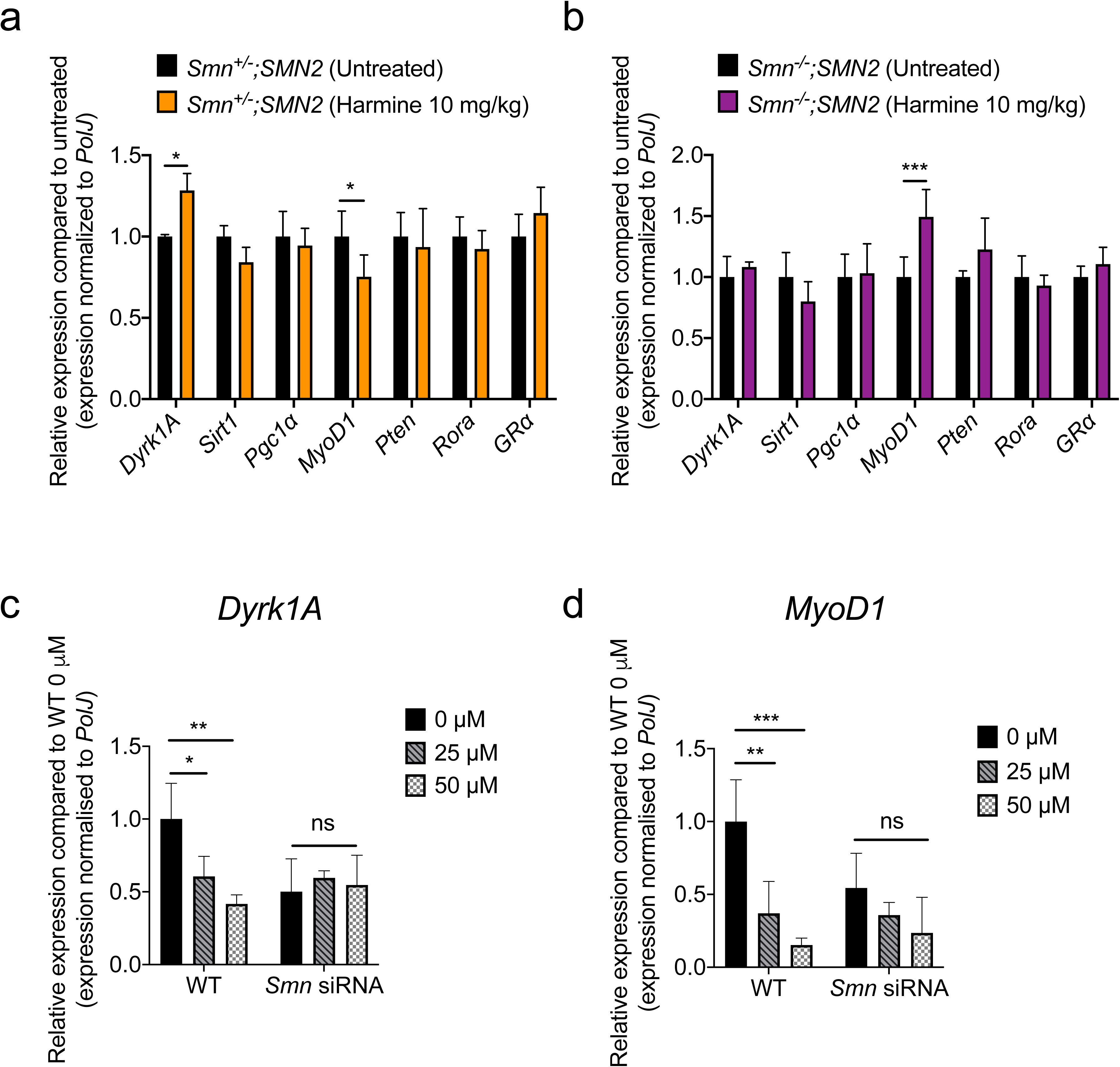
Differential *in vivo* and *in vitro* effects of harmine on muscle expression of predicted molecular effectors. All treated animals received a daily dose of harmine (10 mg/kg, diluted in 0.9% saline) by gavage starting at postnatal day (P) 0. **a.** qPCR analysis of *Dyrk1A*, *Sirt1*, *Pgc1α*, *MyoD1*, *Pten*, *Rora* and *GRα* expression in triceps of P7 untreated and harmine-treated *Smn^+/-^;SMN2* control littermates. Data are mean ± s.d., n = 4 animals for untreated *Smn^+/-^;SMN2* mice and 3 for harmine-treated *Smn^+/-^;SMN2*, two-way ANOVA, \**p*<0.05. **b.** qPCR analysis of *Dyrk1A*, *Sirt1*, *Pgc1α*, *MyoD1*, *Pten*, *Rora* and *GRα* expression in triceps of P7 untreated and harmine-treated *Smn^-/-^;SMN2* mice. Data are mean ± s.d., n = 3 animals for untreated *Smn^-/-^;SMN2* mice and 5 for harmine-treated *Smn^-/-^;SMN2*, two-way ANOVA, \*\*\**p*<0.001. **c.** qPCR analysis *Dyrk1A* expression in differentiated WT and *Smn*-depleted C2C12s that were untreated or exposed to 25 or 50 μM harmine. Data are mean ± s.d., n = 3 for each experimental group, two-way ANOVA, \**p*<0.05, \*\**p*<0.01, ns = not significant. **d.** qPCR analysis *MyoD1* expression in differentiated WT and *Smn*-depleted C2C12s that were untreated or exposed to 25 or 50 μM harmine. Data are mean ± s.d., n = 3 for each experimental group, two-way ANOVA, \*\**p*<0.01, \*\*\**p*<0.001, ns = not significant.

Nevertheless, our bioinformatic analyses uncover several interesting molecular networks restored by harmine in SMA muscle that could have further implications for future development of muscle-specific therapies for SMA.

## DISCUSSION

Despite the tremendous recent advances in SMA gene therapy, this neuromuscular disorder remains incurable and there is an urgent need for the development of second-generation treatments that can be used in combination with SMN-dependent therapies ^17–19^. In this study, we therefore evaluated and validated a strategy combining transcriptomics, proteomics and drug repositioning to identify novel therapeutic compounds that have the potential to improve muscle pathology in SMA. An in-depth investigation of one of these drugs, harmine, further supports our approach as harmine restored several molecular, behavioural and histological disease phenotypes in both cellular and animal models of the disease.

Of major importance, and to our surprise, we demonstrate that early SMN restoration via Pip6-PMO corrects most, if not all, of the transcriptomic and proteomics dysregulations in SMA muscle, highlighting the need for and likely benefit from early treatment intervention in SMA. It is important to note however that the Pip6a-PMO dose delivered to mice was very high and most likely higher than what would be expected in patients. Our pathway analyses reveal that many molecular functions that are dysregulated in SMA mice compared to WT mice and recovered by Pip6a-PMO have previously been implicated in the pathology of SMA such as RNA metabolism and splicing, circadian regulation of gene expression, ubiquitin pathways, regulation of Rho protein signal transduction and actin binding pathways ^44–47^. Their normalization following SMN restoration further supports their involvement in SMA pathology.

Using the differentially expressed genes and protein in SMA muscle compared to WT, we used a CMap pertubational dataset to provide a list of candidate drugs that could improve SMA pathology, some of which had previously been evaluated in SMA such as salbutamol ^48^. CMap analysis has previously been used to identify new potential therapeutics for a range of different conditions such as skeletal muscle atrophy ^49^, osteoarthritic pain ^50^, osteoporosis ^51^, gliomas ^52^, lung adenocarcinoma ^53^, hepatoblastoma ^54^, acute myeloid leukemia ^55^ and kidney disease ^56^. CMap can also help establish prediction models for different adverse drug reactions and to evaluate drug safety ^57^.

In this study, we chose to provide a more in-depth assessment of harmine, a drug predicted to restore differentially expressed genes and proteins in SMA muscle. Interestingly, one of the genes downregulated in SMA muscle and restored by harmine is *Snrnp27*, a small nuclear RNP (snRNP) involved in pre-mRNA splicing ^58^ and SMN plays a canonical role in the assembly of snRNPs ^59^. *Cyclin dependent kinase inhibitor 1A* (*Cdkn1a* or *p21*) was also identified as a potential molecular target of harmine. This mediator of cell cycle and DNA repair is reported to be upregulated in various SMA models ^60–64^. In addition to specific genes, GO analysis of our RNA-Seq data reveals that harmine restores several genes implicated in key muscle processes such as muscle structure development, muscle contraction, muscle system process and muscle cell differentiation. Thus, our combined transcriptomics, proteomics and CMap analysis has identified genes that have previously been implicated in SMA pathology.

While harmine was selected as a non-SMN treatment strategy, we found an upregulation of FL *SMN2* but not total *SMN2* in SMA cells and mice, implying that harmine possibly affects alternative splicing of *SMN2*. Interestingly, harmine restored the alternative splicing of *TNNT2* and *INSR* in DM1 myoblasts and muscle from DM1 mice ^35^, further supporting a role for harmine in modulating mRNA splicing. As demonstrated by our RNA-Seq analysis, harmine further restores the expression of several genes, indicating that its potential benefits may stem from combinatorial effects on SMN and non-SMN genes. The latter most likely makes the greatest contribution to the therapeutic benefits observed, given that harmine did not increase SMN protein levels.

In addition, harmine can cross the blood-brain barrier and has well characterized neuroprotective properties, including its ability to upregulate the expression of GLT-1 in several neurodegenerative models ^40, 41^. We indeed show that GLT-1 expression is reduced in the spinal cord of SMA mice and significantly upregulated following harmine administration. Reduced glutamate transporter activity throughout the CNS of SMA patients has also been reported ^65^. The fact that harmine exerts muscle and CNS effects makes it an interesting therapeutic option for SMA. However, it is important to note that harmine can also exert adverse effects such as the onset of tremors ^37–39^, which we observed when dosing the intermediate *Smn^2B/-^* mouse model over a longer period of time.

To our knowledge, this is the first in-depth validation of this combinatorial approach in SMA. We are able to show strength and potential of combining multi-omics and drug repositioning to uncover novel therapeutic entities, which in this case was aimed at improving muscle health in SMA. Our work thus provides an invaluable list of pharmacological compounds that can be evaluated for treatment of SMA muscle pathology as well as strong support for the use of this combined multi-omics and bioinformatic strategy.

## ONLINE METHODS

### Animals and animal procedures

Wild-type mice (FVB/N ^66^ and C57BL/6J ^67^) were obtained from Jackson Laboratories. The severe *Smn^-/-^;SMN2^+/-^* mouse model ^68^ was also obtained from Jackson Laboratories (FVB.Cg-Smn1tm1Hung Tg(SMN2)2Hung/J). The moderate *Smn^2B/-^* mouse model ^69^ was generously provided by Dr. Lyndsay M Murray, University of Edinburgh). All experiments with live animals were performed at the Biomedical Services Building, University of Oxford. Experimental procedures were authorized and approved by the University of Oxford ethics committee and UK Home Office (current project license PDFEDC6F0, previous project license 30/2907) in accordance with the Animals (Scientific Procedures) Act 1986.

The Pip6a-PMO and Pip6a-scrambled conjugates were both separately prepared in 0.9% saline solution and administered at a dose of 10 µg/g via an intravenous facial vein injection at P0 and P2.

Harmine hydrochloride (sc-295136, Insight Biotechnology Ltd, Sante Cruz) was dissolved in 0.9% saline and administered daily (10 mg/kg) by gavage.

### Synthesis of Pip6a peptide-PMO conjugates

The PMO sequence targeting ISS-N1 intron 7 (−10-27) (5′-ATTCACTTTCATAATGCTGG-3′) and scrambled PMO (5’-TAC GTT ATA TCT CGT GAT AC-3’) were purchased from Gene Tools LLC (Corvallis).

The Pip6a Ac-(RXRRBRRXRYQFLIRXRBRXRB)-COOH peptide was manufactured by standard 9-fluorenylmethoxy carbonyl chemistry, purified to >90% purity by reverse-phase high-performance liquid chromatography (HPLC) and conjugated to the 3’ end of the PMO through an amide linkage. The conjugate was purified by cation exchange HPLC, desalted and analyzed by mass spectrometry. Pip6a peptide-PMO conjugates were dissolved in sterile water and filtered through a 0.22 µm cellulose acetate membrane before use.

### Laminin staining of skeletal muscle

*Tibialis anterior* (TA) muscles were fixed in 4% PFA overnight. Tissues were sectioned (13 μm) and incubated in blocking buffer for 2 hours (0.3% Triton-X, 20% FBS and 20% normal goat serum in PBS). After blocking, tissues were stained overnight at 4 °C with rat anti-laminin (Sigma) in blocking buffer. The next day, tissues were washed in PBS and probed using goat-anti-rat IgG 488 secondary antibodies (Invitrogen) for one hour. PBS-washed tissues were mounted in Fluoromount-G (Southern Biotech). Images were taken with a DM IRB microscope (Leica) with a 20× objective. Quantitative assays were performed blinded on 3–5 mice for each group and five sections per mouse. The area of muscle fiber within designated regions of the TA muscle sections was measured using Fiji ^70^.

### qPCR

RNA was extracted from tissues and cells by either an RNeasy kit from Qiagen or by guanidinium thiocyantate-acid-phenol-chloroform extraction using TRIzol Reagent (Life Technologies) as per manufacturer’s instructions. The same RNA extraction method was employed for similar experiments and equal RNA amounts were used between samples within the same experiments. cDNA was prepared with the High Capacity cDNA Kit (Life Technologies) according to the manufacturer’s instructions. The cDNA template was amplified on a StepOnePlus Real-Time PCR Thermocycler (Life Technologies) with SYBR Green Mastermix from Applied Biosystems. qPCR data was analyzed using the StepOne Software v2.3 (Applied Biosystems). Primers used for qPCR were obtained from IDT and sequences for primers were either self-designed or ready-made (Supplementary Table 2). Relative gene expression was quantified using the Pfaffl method ^71^ and primer efficiencies were calculated with the LinRegPCR software. We normalized relative expression level of all tested genes in mouse tissue and cells to *RNA polymerase II polypeptide J* (*PolJ*) ^72^. For human cells, we ran a GeNorm kit (Primer Design) to identify *RPL13A* as a reference/housekeeping gene. Primers for *RPL13A* were from IDT (209604333).

### Cell culture

Both C2C12s ^73^ and NSC-34s ^74^ cell lines were maintained in growth media consisting of Dulbecco’s Modified Eagle’s Media (DMEM) supplemented with 10% fetal bovine serum (FBS) and 1% Penicillin/Streptomycin (all Life Technologies). The cells were cultured at 37°C with 5% CO^2^. C2C12 myoblasts were differentiated in DMEM containing 2% horse serum (HS) for 7 days to form multinucleated myotubes.

For siRNA experiments, C2C12 cells were seeded in 12-well plates and after reaching 50% confluence, growth media was changed to differentiation media and the cells were transfected with 10 µM of siSmn (Duplex name: mm.RiSmn1.13.1) and scrambled siRNA (scrambled negative control DsiRNA, #51-01-19-08) (both from IDT) in an siRNA-lipofectamine complex (Lipofectamine® RNAiMAX Reagent, Life Technologies). Fresh media containing the transfection reagents was changed every 2 days. At D6, the C2C12 myotubes were further exposed to harmine (25 and 50 µM) for 48 hours.

Human fibroblasts were obtained from Coriell Institue (SMA GM03813, control AG02261) and cultured in DMEM, supplemented with 1% antibiotics/antimycotics and 20% FBS.

### MTS assays

Cell viability and proliferation of C2C12 and NSC-34 cells treated with harmine (sc-202644, Insight Biotechnology Ltd, Sante Cruz) dissolved in DMSO (final concentration 0.03%) were evaluated with a 3-(4,5-dimethylthiazol-2-yl)-5-(3-carboxymethoxyphenyl)-2-(4-sulfophenyl)-2H-tetrazolium (MTS) assay kit (Colorimetric). The measurements were made according to manufacturer’s instructions. Briefly, 10 µl of MTS reagent was added directly to the wells and cell plates were incubated at 37°C for a minimum of 1 hour. Absorbance was measured at 490 nm on a CLARIOstar® plate reader (BMG LABTECH). Background absorbance was first subtracted using a set of wells containing medium only, then normalized to and expressed as a relative percentage of the plate-averaged untreated control. To chemically induce apoptosis, cells were treated with 10 μM Staurosporine (Abcam, Cambridge, UK).

### Western blot

Freshly prepared radioimmunoprecipitation (RIPA) buffer was used to homogenize tissue and cells, consisting of 50 mM Tris pH 8.8, 150mM NaCl, 1% NP-40, 0.5% Sodium Deoxycholate, 0.1% SDS and complete mini-proteinase inhibitors (1 tablet per 10 ml extraction solution, Roche). Equal amounts of total protein were loaded, as measured by Bradford Assay. Protein samples were first diluted 1:1 with Laemmli sample buffer (Bio-Rad, Hemel Hempstead, UK) containing 5% β-mercaptoethanol (Sigma) and heated at 100°C for 10 minutes. Next, samples were loaded on freshly made 1.5 mm 12% polyacrylamide separating and 5% stacking gel and electrophoresis was performed at 120 V for ∼1.5h in running buffer. Subsequently, proteins were transferred from the gel onto to a polyvinylidene fluoride (PVDF) membrane (Merck Millipore) via electroblotting at 120 V for 60 minutes in transfer buffer containing 20% methanol. Membranes were then incubated for 2h in Odyssey Blocking Buffer (Licor). The membrane was then probed overnight at 4°C with primary antibodies (anti-GLT-1, 1:1000, Abcam #ab41621; anti-SMN, 1:1000, Millipore #MABE230; anti-vinculin, 1:200.000, Sigma #V9131) in Odyssey Blocking Buffer and 0.1% Tween-20. The next day, after three 10-minute washing steps with PBS, the membrane was incubated for 1 hour at room temperature with secondary antibodies conjugated to infrared dyes. Lastly, the membrane was washed again three times 10 minutes in PBS and visualized by scanning 700 nm and 800 nm channels on the LI-COR Odyssey CLx infrared imaging system (LI-COR) for 2.5 minutes per channel. The background was subtracted and signal of protein of interest was divided by signal of the housekeeping protein or total protein, per sample.

### Proteomic analysis

Proteomic analyses were performed using a liquid chromatography–mass spectrometry (LC-MS)-based method. High-resolution isoelectric focusing (HiRIEF) was used at the peptide level in the 3.7–5.0 pH range. Two tandem mass tags (TMTs, chemical labels) were used for mass spectrometry (MS)-based quantification and identification of proteins. The data was median normalized based on peptide ratio. Amongst a total of 9798 potentially detectable proteins, most (8152) were identified in all samples/groups.

The limma R package was used for differential expression analysis, whereby differentially expressed proteins were defined by FDR < 0.05. Gene Ontology enrichment analysis of proteomic data was executed using topGO R function and adjusted *p* values for multiple testing following a Benjamini-Hochberg correction. For principal component analysis, we used the prcomp R function on the normalized expression data.

### Microarray analysis

RNA was extracted by guanidinium thiocyantate-acid-phenol-chloroform extraction using TRIzol Reagent (Life Technologies) as per manufacturer’s instructions. GeneChip Mouse Transcriptome Assay 1.0 arrays were used (Affymetrix core facility, Karolinska Institute) with 100 ng of RNA per sample. Annotations for the Mouse Transcript Array 1.0 at the transcript level were obtained from the Affymetrix website (http://www.affymetrix.com/products_services/arrays/specific/mo_trans_assay. affx#1_4). We performed background correction and RMA normalization at the probe level using oligo R package. We summarized the data in ensemble transcript IDs using the average. The total number of ensemble transcript IDs was 93,594, corresponding to 37,450 genes. For differential expression analysis, we used limma R package and considered a transcript differentially expressed if their FDR < 0.05. A gene was considered differentially expressed if at least one of the associated transcripts was differentially expressed. Gene Ontology enrichment analysis was performed in R using the topGO function as described for proteomic data. For principal component analysis we used the prcomp R function on the RMA normalized gene expression data at the gene level (for comparison with proteomic data).

### Combined analysis of proteomic and transcriptomic data

To measure the similarity between gene expression profiles, we used the Ward hierarchical clustering on the Euclidean distance of 1–r (where r is the Pearson correlation between samples). To compare the two omics readouts, proteomic and transcriptomic data were scaled (transformed to z-score values), followed by a PCA analysis showing that PC1 divides the data at the transcript and protein level (Supplementary Fig. 1). Using the kill.pc function in the swamp R package, we extracted a new expression matrix where the variance given by PC1 has been removed. Finally, we performed hierarchical clustering analysis on the new expression matrix.

### RNA-Sequencing analysis

RNA was extracted using a RNeasy Microarray Tissue Mini Kit from Qiagen. Lysis and homogenization were performed using QIAzol Lysis Reagent. cDNA synthesis and RNA-Seq library construction were performed at the Oxford Genomics Centre (Oxford, United Kingdom) using poly(A) enrichment of the mRNA (mRNA-Seq) and HiSeq 4000 Systems for sequencing. All samples passed quality control. For differential expression analysis, we used DESeq2 on genes expressed across all samples (15523 genes) after removal of one outlier (Harmine-treated SMA sample 1). We considered a gene differentially expressed at FDR < 0.05. For Gene Ontology enrichment analysis, we used topGO R function and adjusted p values for multiple testing following a Benjamini-Hochberg correction. For mouse phenotype enrichment analysis, we downloaded phenotypes from the Mouse Genome Database (MGD), Mouse Genome Informatics, The Jackson Laboratory, Bar Harbor, Maine (URL: http://www.informatics.jax.org) (June, 2018) and used in-house script to correct for the background set of expressed genes.

### Gene functional network and clustering method

A gene functional network is built by extracting interactions from a phenotypic linkage network ^43^ for the top 500 differentially expressed genes between WT and SMA samples. To identify modules of highly interconnected genes in the network, we employed “cluster_louvain” function in “igraph” R package ^75^. This function implements the multi-level modularity optimization algorithm ^76, 77^ where at each step genes are re-assigned to modules in a greedy way and the process stops when the modularity does not increase in a successive step.

### Upstream regulators

Ingenuity Pathway Analysis (www.qiagenbioinformatics.com) was used to identify the top 100 upstream regulators for the top 500 differentially expressed genes between WT and SMA samples. A reduced list of regulators was identified based on their target genes to be within the four harmine-reversed modules.

### CMap analysis

Ensembl transcript ids from mice were mapped to human probe IDs (HG-U133A) using biomaRt (Ensembl transcript id *mus musculus* → Ensembl gene id *mus musculus* → ortholog_one2one → Ensembl gene id *homo sapiens* → HG-U133A id). We compared the identified disease and Pip6a-PMO signatures (top 500 up-regulated and top 500 down-regulated genes/proteins) to the drug instances contained in the CMap dataset (Build 02, http://www.broadinstitute.org/cmap), which are defined as the basic unit of data and metadata in CMap.

### Statistical Analysis

All statistical analyses were done with the most up-to-date Graphpad Prism software. When appropriate, a Student’s unpaired two-tail *t*-test, a one-way ANOVA followed by a Tukey’s multiple comparison test or a two-way ANOVA followed by a Sidak’s multiple comparison test was used. Outliers were identified via the Grubbs’ test. For the Kaplan-Meier survival analysis, the log-rank test was used and survival curves were considered significantly different at *p*<0.05.

## SUPPLEMENTARY FIGURE LEGENDS

**Supplementary Figure 1. Clustering and principal component analysis integrating proteomic and transcriptomic data. a.** Heatmap correspond to the Pearson correlation between each pair of proteomic and transcriptomic profiles. Gene expression profiles show higher correlations than proteomic data, but similar clustering of experimental groups is observed within each type of data. **b.** Principal component analysis of the proteomic and transcriptomic data separates by data by type (RNA or Protein) along the first component.

**Supplementary Figure 2.** Principal component analysis on proteomic (**a**) and transcriptomic (**b**) data of untreated post-natal day (P) 2 and P7 *Smn^-/-^;SMN2* and WT mice.

**Supplementary Figure 3. *In vitro* Dose-dependent toxicity of harmine treatment *in vitro*.** C2C12s (a) and NSC-34s (b) were treated with 1, 10, 25 or 50 µM for 24, 48 or 72 hours. Control groups were untreated cells or cells treated with either DMSO (vehicle) or Staurosporine (positive control). An MTS assay was performed on all experimental groups and MTS scores are normalized to untreated cells at 24 hours (100%). Data are mean ± s.d., n = 3 independent wells, two-way ANOVA, \*\**p*<0.01, \*\*\**p*<0.001, \*\*\*\**p*<0.0001.

## SUPPLEMENTARY TABLES

**Supplementary Table 1.** Proteins downregulated in P7 Pip6a-PMO-treated *Smn^-^*^/-^;SMN2 mice compared to P7 untreated WT mice.

**Supplementary Table 2.** List of mouse and human qPCR primers.

## SUPPLEMENTARY FILES

**Supplementary File 1.** Top biological processes enriched among the genes differentially expressed in disease and Pip6a-PMO treatment.

## ACKNOWLEDGEMENTS

We would like to thank the staff at the BMS facility at the University of Oxford, Dr Emelie Blomberg and Dr Samir El-Andaloussi (Karolinska Institute) for the microarray services, Dr Henrik Johansson (Karolinska Institute) for the proteomic services and the Oxford Genomics Centres for the RNA Sequencing services. K.E.M. was funded by the MDUK and SMA Trust. M.B. was funded by the SMA Trust. J.M.H. is funded by the Keele University School of Medicine. S.M.H. is funded by the MRC DPFS (MR/R025312/1). Computation used the Oxford Biomedical Research Computing (BMRC) facility, a joint development between the Wellcome Centre for Human Genetics and the Big Data Institute supported by Health Data Research UK and the NIHR Oxford Biomedical Research Centre. The views expressed are those of the author(s) and not necessarily those of the NHS, the NIHR or the Department of Health.

