## Supplementary figures and images for "Combining multi-omics and drug perturbation profiles to identify novel treatments that improve disease phenotypes in spinal muscular atrophy"

### Supplementary Figure 1

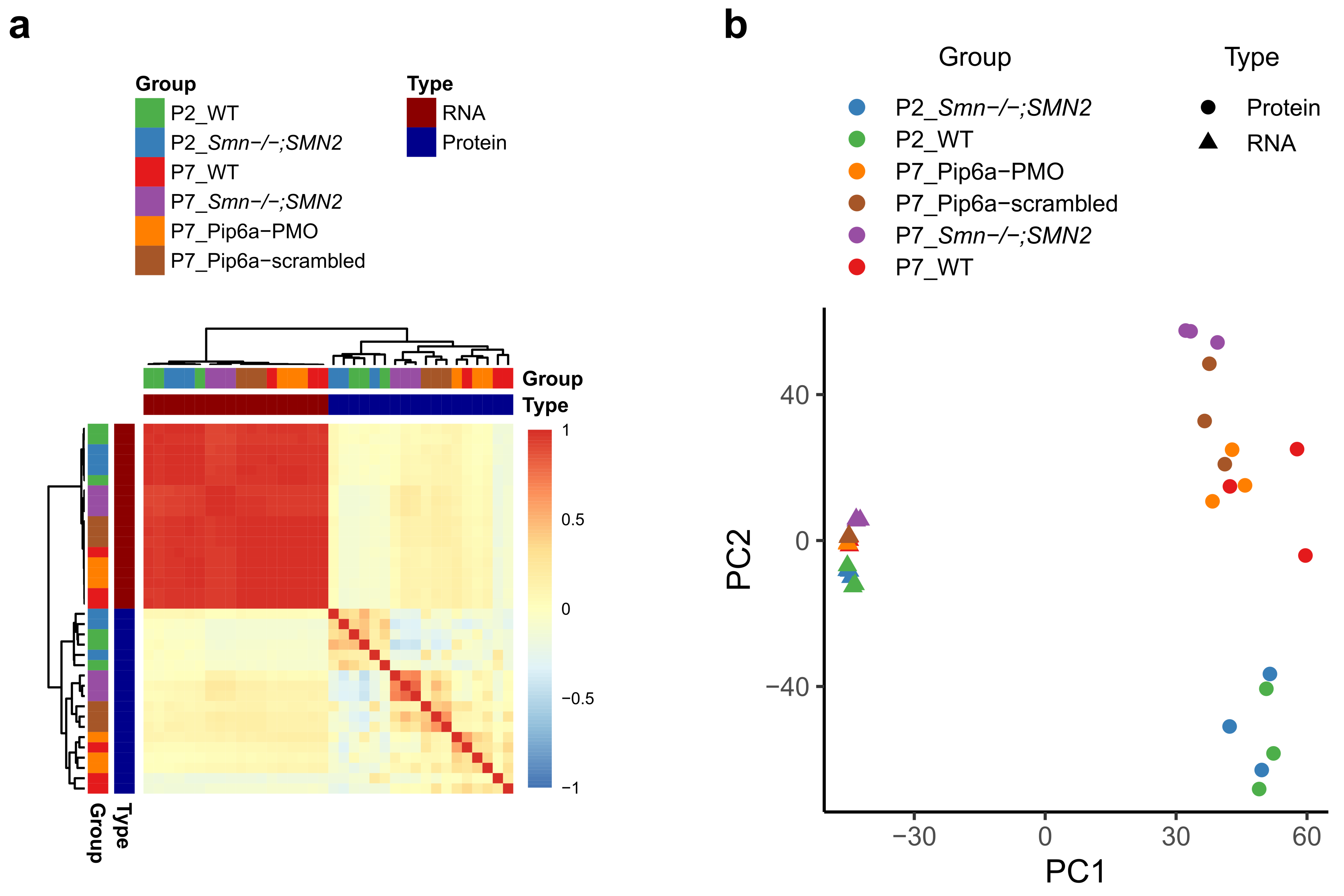

### Supplementary Figure 2

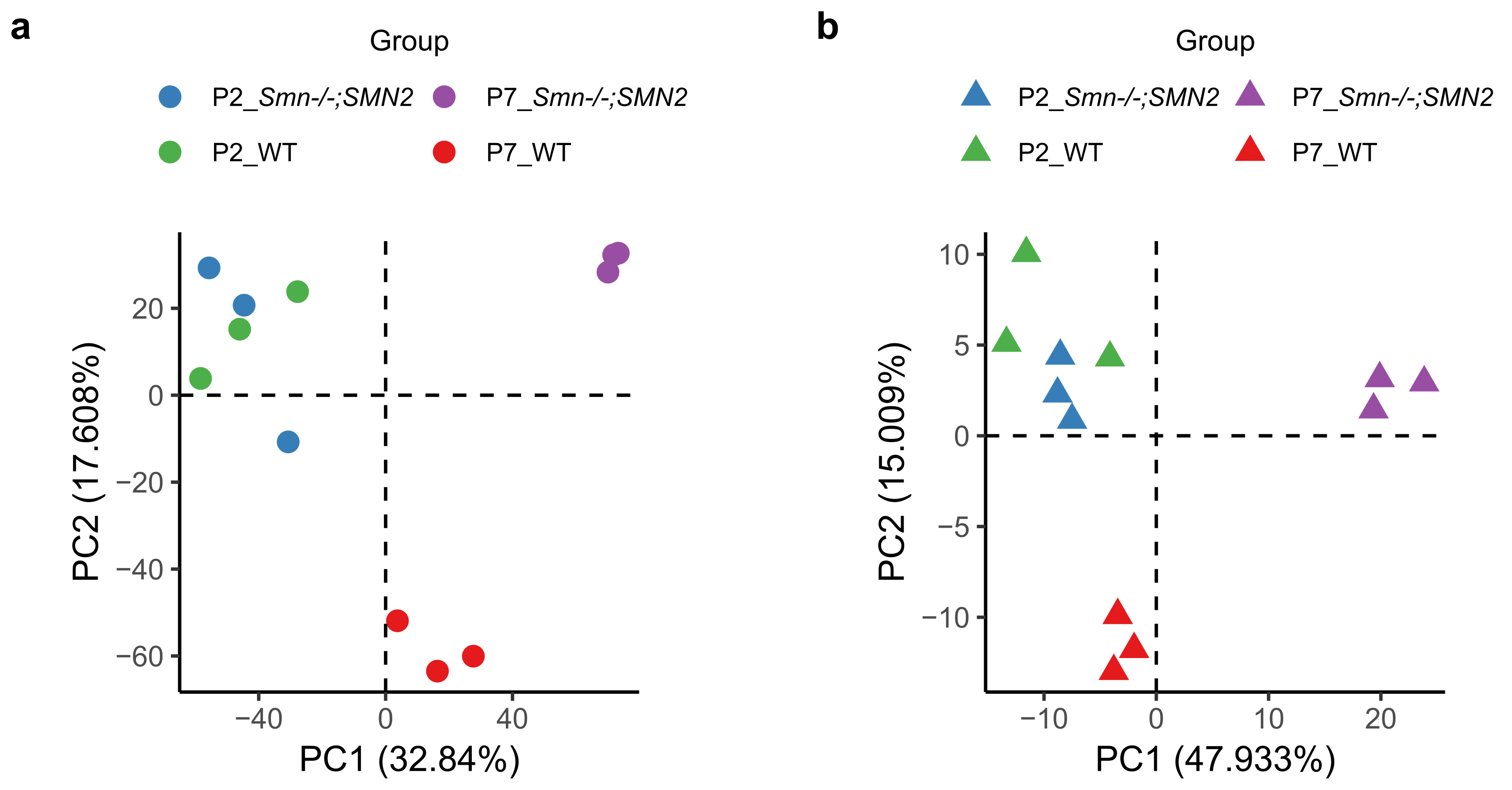

### Supplementary Figure 3

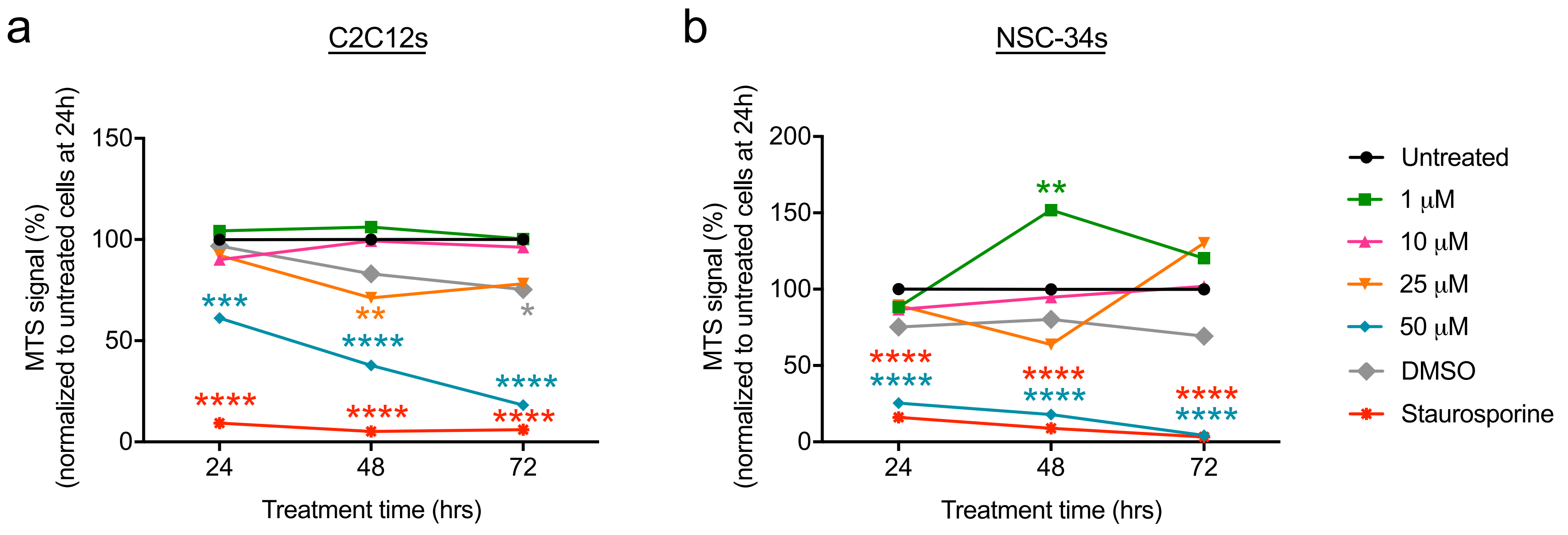
