## Supplementary Table 1 for "Combining multi-omics and drug perturbation profiles to identify novel treatments that improve disease phenotypes in spinal muscular atrophy"

**Proteins downregulated in Pip6a-PMO treated SMA mice vs. WT mice**

Immunoglobulin kappa variable 4-53

Immunoglobulin kappa variable 10-95

Immunoglobulin kappa variable 8-28

Immunoglobulin kappa variable 8-19

Immunoglobulin kappa constant

Immunoglobulin heavy constant gamma 1

Immunoglobulin heavy variable 3-5

Immunoglobulin heavy variable 1-81

TAP binding protein

Tap1 transporter 1, ATP-binding cassette, sub-family B

Survival Motor Neuron

* Proteins were considered downregulated if false discovery rate (FDR) < 0.05.
