## Supplementary Table 2 for "Combining multi-omics and drug perturbation profiles to identify novel treatments that improve disease phenotypes in spinal muscular atrophy"

| Mouse | Forward | Reverse |
| --- | --- | --- |
| *Aspm* | 5′-TTCTATGACGAACGCTGGAAG-3' | 5′-CTTTCCGCTCCCAAAACAAG-3' |
| *Cdkn1a* | 5′-CAGATCCACAGCGATATCCAG-3' | 5′-AGAGACAACGGCACACTTTG-3' |
| *Clpx* | 5′-ACAAATACTGACCGAGCCAC-3' | 5′-TTCTTTCCAGGGCCAATCTC-3' |
| *Gls* | 5′-GTGGTTTCTGCCCAATTACTG-3' | 5′-CCCAGCAACTCCAGATTTTG-3' |
| *Mcm2* | 5′-AAAGCATCTCCATCTCCAAGG-3' | 5′-GGCTCTGTGAGGTCTACATTC-3' |
| *Ppm1b* | 5′-CTTTCTACCTTCGCCCCAG-3' | 5′-ATCTTCCATTTCTACTCTCCATCC-3' |
| *SnrnP27* | 5′-CTGGAGGGTAAAACAGAGGAAG-3' | 5′-AGACACGTTGATGGCATAGG-3' |
| *Tob2* | 5′-CCTCAGCTACAACCTGAATACC-3' | 5′-TCTTTCCCTCTTGCGTTTGG-3' |
| *Smn* | 5’-TGCTCCGTGGACCTCATTTCTT-3’ | 5’-TGGCTTTCCTGGTCCTAATCCTGA-3’ |
| *Dyrk1a* | 5’-GAGACACACAGTCCCCAGGT-3’ | 5’-ACTGTGGCCAACCTCCATAG-3’ |
| *Sirt1* | 5’-ACGCTGTGGCAGATTGTTAT-3’ | 5’-GCAAGGCGAGCATAGATA-3’ |
| *Pgc1a* | 5’-TGGAGTGACATAGAGTGTGCTGC-3’ | 5’-CTCAAATATGTTCGCAGGCTCA-3’ |
| *MyoD1* | 5’-ATCCGCTACATCGAAGGTCT-3’ | 5’-CGCTGTAATCCATCATGCCA-3’ |
| *Pten* | 5’-CATAACCCACCACAGCTAG-3’ | 5’-GCAGACCACAAACTGAGG-3’ |
| *Rora* | 5’-TGCGAGCTCCAGCCGAGGTA-3’ | 5’-GCCCTTGCAGCCTTCACACGTA-3’ |
| *GRα* | 5’-AAAGAGCTAGGAAAAGCCATTGTC-3’ | 5’-TCAGCTAACATCTCTGGGAATTCA-3’ |

| **Human** | **Forward** | **Reverse** |
| --- | --- | --- |
| *ASPM* | 5′-GGACAAAACCCATTATCGCTG-3' | 5′-CCCTGTTCCTGCTTTTCCTTC-3' |
| *CDKN1a* | 5′-TGTCACTGTCTTGTACCCTTG-3' | 5′-GGCGTTTGGAGTGGTAGAA-3' |
| *CLPX* | 5′-GTGCTTTCAGTTGCTGTGTAC-3' | 5′-TCATCCTCCCGTCTTCTTATTTC-3' |
| *GLS* | 5′-TTCCAGAAGGCACAGACATG-3' | 5′-GGCTCAGTACTCTTTCACCAG-3' |
| *MCM2* | 5′-ATTTCGTCCTGGGTCCTTTC-3' | 5′-CGCTGGTAGTTCTGATAGATGG-3' |
| *Ppm1b* | 5′-CCCTTGCCTCAGATTTATTGC-3' | 5′-TTCCATTTCCACTCTCCATCC-3' |
| *SMN, full-length* | 5′-GCTTTGGGAAGTATGTTAATTTCA-3' | 5′-CTATGCCAGCATTTCTCCTTAATT-3' |
| *SMN, total* | 5′-GCGATGATTCTGACATTTGG-3' | 5′-GGAAGCTGCAGTATTCTTCT-3' |
| *SnRNP27* | 5′-AGAAACAAAGAGCAAAGAACGG-3' | 5′-TTTACAGAGCCATCCACCTTC-3' |
| *Tob2* | 5′-AGCTACAACCTGAACACCATG-3' | 5′-TCTCTTTTCTGTGGTCTTGGG-3' |
